# Germline Contamination and Leakage in Whole Genome Somatic Single Nucleotide Variant Detection

**DOI:** 10.1101/204370

**Authors:** Dorota H. Sendorek, Cristian Caloian, Kyle Ellrott, J. Christopher Bare, Takafumi N. Yamaguchi, Adam D. Ewing, Kathleen E. Houlahan, Thea C. Norman, Adam A. Margolin, Joshua M. Stuart, Paul C. Boutros

## Abstract

**Background:** The clinical sequencing of cancer genomes to personalize therapy is becoming routine across the world. However, concerns over patient re-identification from these data lead to questions about how tightly access should be controlled. It is not thought to be possible to re-identify patients from somatic variant data. However, somatic variant detection pipelines can mistakenly identify germline variants as somatic ones, a process called “germline leakage”. The rate of germline leakage across different somatic variant detection pipelines is not well-understood, and it is uncertain whether or not somatic variant calls should be considered re-identifiable. To fill this gap, we quantified germline leakage across 259 sets of whole-genome somatic single nucleotide variant (SNVs) predictions made by 21 teams as part of the ICGC-TCGA DREAM Somatic Mutation Calling Challenge.

**Results:** The median somatic SNV prediction set contained 4,325 somatic SNVs and leaked one germline polymorphism. The level of germline leakage was inversely correlated with somatic SNV prediction accuracy and positively correlated with the amount of infiltrating normal cells. The specific germline variants leaked differed by tumour and algorithm. To aid in quantitation and correction of leakage, we created a tool, called GermlineFilter, for use in public-facing somatic SNV databases.

**Conclusions:** The potential for patient re-identification from leaked germline variants in somatic SNV predictions has led to divergent open data access policies, based on different assessments of the risks. Indeed, a single, well-publicized re-identification event could reshape public perceptions of the values of genomic data sharing. We find that modern somatic SNV prediction pipelines have low germline-leakage rates, which can be further reduced, especially for cloud-sharing, using pre-filtering software.

## Background

The appropriate limits on data sharing remains a contentious issue throughout biomedical research, as shown by recent controversies [1]. Studies such as the Personal Genome Project (PGP) have pioneered open sharing of patient data for biomedical research, while ensuring that enrolled patients consent to risks of identification [2]. In fact, analysis of PGP data has showed that a majority of participants can be linked to a specific named individual [3]. Identifiability is greatly facilitated when researchers release all generated data online – as is standard in some fields [4]. This public, barrier-free release has numerous advantages. It can minimize storage costs, increase data redundancy to reduce the risk of data-loss and maximize data availability and re-use. As a result, it is argued that barrier-free deposition of genomic data in public repositories like GEO [5-6] or dbGaP [7-8] promotes collaborative work and maximizes the value of already-funded research [9]. Further, many researchers believe they have an ethical duty to release all data [10].

Nevertheless, there are at least four counter-arguments in favour of a conservative approach to data protection. First, the groups generating the data have uniquely intimate knowledge of it and studies done without their participation can be more prone to errors, although improved documentation of the research process can mitigate this effect [1]. Second, the desire to immediately release data may oppose the desire to explore complex inter-linked questions. The initial report of a dataset may not fully reflect the magnitude of work that goes into generating it, particularly for clinical trials. With immediate data release, the data collectors may find themselves under time constraints, unable to comprehensively exploit the data they produced without competition from subsequent researchers who are able to use the data freely. This effectively disincentivizes the challenging work of dataset creation, producing a situation akin to a tragedy of the commons. Third, the inherent value in large datasets may enable data producers to seek commercialization opportunities by keeping data resources private. Fourth, many studies involve data derived from human subjects that contain revealing and personal information, which is under legal protection [11]. Legislation designed to protect patient privacy, such as the Health Insurance Portability and Accountability Act (HIPAA) [12], the Common Rule [13] and the European Union’s General Data Protection Regulation [14] impose harsh financial and professional penalties for violations. As genomic data becomes widely available and techniques for interpreting them improve, de-identification grows increasingly difficult, challenging implementation of barrier-free access that upholds ethical considerations. We focus here on this fourth challenge, or re-identifiability.

Earlier studies have quantified how much DNA information is required to identify individuals. One suggests that as few as 30-80 statistically-independent single nucleotide polymorphisms (SNPs) suffice [15]. Under certain circumstances, small segments of DNA can even be used to recover participants’ names by accessing publicly available, commercial genealogy websites [16]. These problems are compounded by deficiencies in techniques used to prevent re-identification: for example, pooling DNA samples does not prevent detection of any individual sequence [17]. More recently, research into information leakage demonstrated how easily patients can be linked back to data from which they previously had been disassociated by correlating seemingly disparate features, namely from phenotypic and genotypic datasets, in what is referred to as a ‘linking attack’ [18-19].

In cancer research, many studies concentrate on identifying somatic mutations that are induced in the process of tumourigenesis and tumour evolution. Identifying these causative mutations can lead to discovery of novel biomarkers and potential therapeutic targets, making public data release critical for accelerating research. Because these mutations are found in the tumour and not in an individual’s germline genome, they do not, by themselves, provide identifying information. Barrier-free release of somatic mutational data can, in theory, occur without compromising patient privacy.

However, tools used to distinguish somatic mutations from germline are imperfect, and sometimes the predicted somatic mutations are in fact germline genetic variants. This “germline leakage” can occur in several ways. Most next-generation sequencing (NGS) base calling algorithms have low error rates [20], including both undetected true variants (false negatives) while some non-existent variants get reported (false positives). These false positives can occur for several reasons, including low coverage (number of reads aligning to a specific position in the genome), which reduces statistical confidence [21]. Even datasets with high total coverage have variable coverage across the genome with particular regions getting sampled at lower rates either through stochastic or structurally biased factors. As a result, sets of somatic variant predictions can be contaminated with germline variants, particularly in the case of single nucleotide variants (SNVs). To account for these errors, some groups filter out any variant seen in a germline database like dbSNP, while others allow only release of mutations in the exome [22]. Still others allow public release of somatic variant predictions from the whole genome [23]. These variations reflect differing views on the likelihoods and risks of germline leakage, and many groups have not yet developed or articulated specific policies.

To help improve our understanding of the magnitude of germline leakage, we analyzed a set of 259 somatic mutation predictions made by 21 groups from around the world on three synthetic tumours during the ICGC-TCGA DREAM Somatic Mutation Calling-DNA (SMC-DNA) Challenge [24]. We developed a software tool, called GermlineFilter, which can help to quantify and mitigate the risks of germline leakage for publicly available somatic SNV data.

## Results

### Gold Standards of Germline Leakage

We sought to evaluate the extent of germline contamination in contemporary cancer whole-genome sequencing (WGS) datasets, particularly those comprising somatic SNV predictions across the entire genome. To do so, we exploited the synthetic data from the ICGC-TCGA DREAM SMC-DNA Challenge [24-25], which benchmarked somatic SNV predictions using synthetic and real tumour-normal whole-genome pairs/. The generation of the synthetic tumours and their properties are fully detailed in Ewing *et al.* [25]. Briefly, high coverage binary alignment map (BAM) files were obtained from cell lines HCC1143 and HCC1954 [26]. BAMSurgeon [25] was used to randomly ‘spike-in’ germline mutations into the BAM files. Each file was then split into two: one file representing a synthetic tumour and the other file representing the matched normal. The tumour BAM file was finalized by adding somatic mutations: both SNVs and structural variants. This methodology allows for the creation of a “gold standard” dataset in which the precise locations of germline and somatic variants are known, enabling comprehensive assessment of leaked germline mutations. We focused on the first three synthetic tumours from SMC-DNA, referred to as IS1, IS2 and IS3. These tumours vary in the number of mutations, normal contamination and subclonal complexity (**Additional file 1: Supplementary Table 1**) [25]. The synthetic tumours have been available to the public for several years and have thus accumulated a large number of somatic mutation calling results from various submitted methods. Additionally, the organizers ran several widely used algorithms with default settings as a baseline [25]. In total, we evaluated 5,792,868 somatic mutations that included 259 analyses by 21 teams across the three tumours (n_IS1_ = 120; n_IS2_ = 71; n_IS3_ = 68).

### Assessment of Germline Leakage

To quantify germline leakage in submissions to the SMC-DNA tumours, we created a Python program called GermlineFilter, which simultaneously evaluates germline leakage in somatic SNV predictions and filters them in real-time to allow barrier-free access to the final results. The overall process has two steps (Figure 1). During the initial preprocessing step, a germline caller is run on paired tumour and normal BAM files to generate the germline variant calls. Current germline callers have high accuracy rates which can be attributed to diploidy-based assumptions of normal human tissue, assumptions that do not hold for somatic variants due to a host of issues (*e.g*. intra-tumour heterogeneity, tissue cellularity, genomic instability). In the following step, each germline SNP is compared against the somatic SNV predictions to be filtered, provided in standard variant call format (VCF), and the matches are identified. Finally, somatic SNV calls can now be filtered, either by rejecting entire submissions that exceed an acceptable level of leakage or by simply removing the calls that match a germline variant.

**Figure 1:**
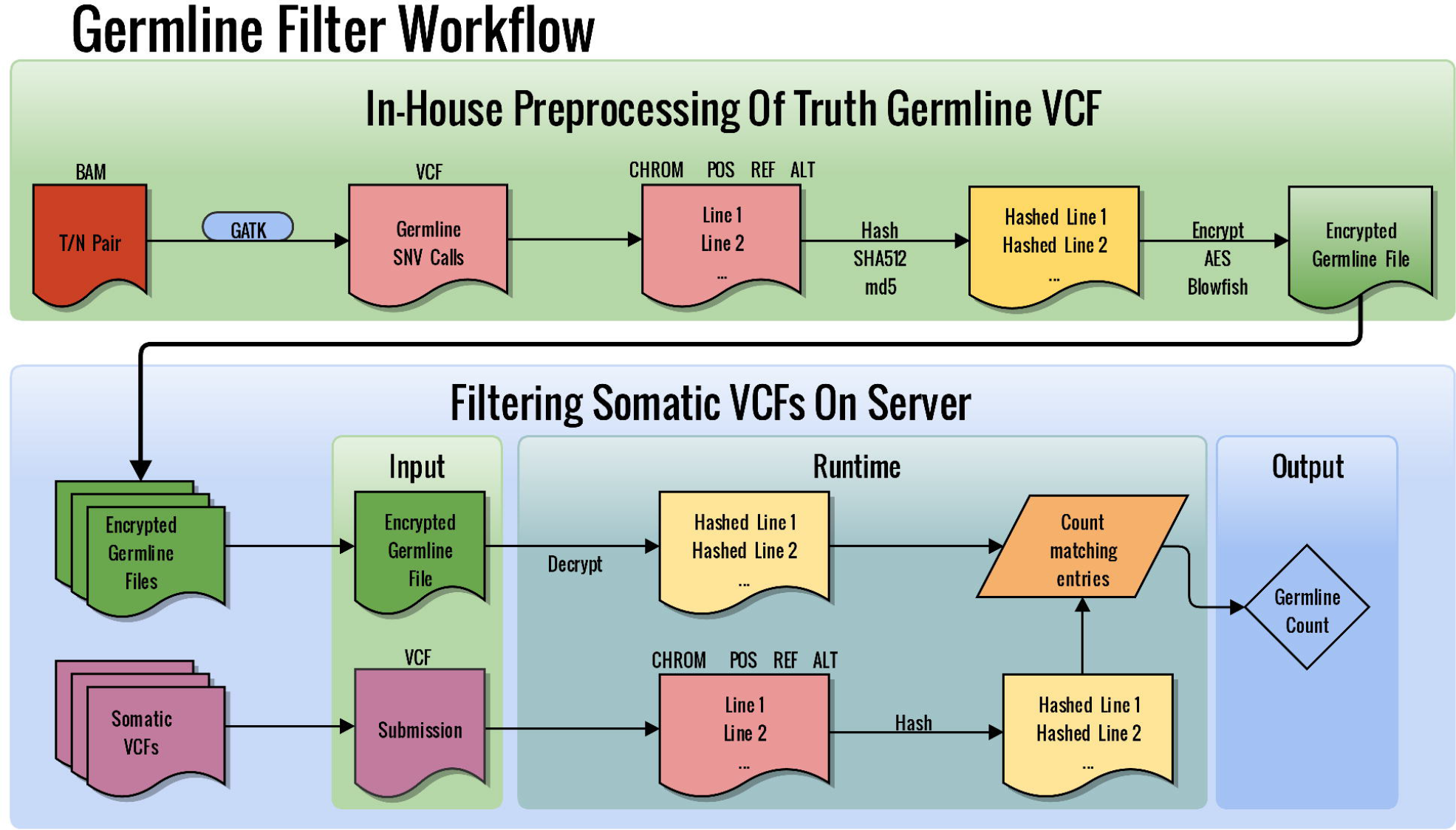
GermlineFilter Workflow. Locally, tumour-normal BAM files are submitted to a germline caller (*e.g*. GATK) to create a germline SNV call VCF file, which is later hashed and encrypted. The encrypted, hashed germline calls can now be moved to any server and used to filter for germline leakage in somatic SNV call VCF files. The output is the germline count found in the somatic calls.

### Germline Contamination Reduces Somatic SNV Prediction Accuracy

The 259 somatic call VCFs submitted during the IS1, IS2 and IS3 phases of the SMC-DNA challenge contained a median of 4,325 SNV calls (averaging 22,366 SNV calls). Each of these was run through GermlineFilter to quantify germline leakage in terms of the number of true germline SNPs misidentified as somatic SNVs. Prediction accuracy for each submission was measured using the F_1_-score (*i.e.* the harmonic mean of precision and recall) in keeping with the metrics used in the DREAM SMC-DNA challenge.

Germline leakage was highly variable across submissions, ranging from 0 to 45,300, with a median of 1 per submission. The median leakage rate across tumours ranged from 0 (IS3), to 2 (IS 1) and went up as high as 6 (IS2). IS2 contained the highest normal contamination (20%), suggesting that even low normal contamination can increase germline leakage. For each tumour, we compared germline count to the previously reported F1-scores (Figure 2a) and found a highly significant negative correlation in each of the three tumours (Spearman’s *ρ*_IS1_ = −0.557, *ρ*_IS2_ = −0.477, *ρ*_IS3_ = −0.410, **Additional file 1: Supplementary Table 1**). For a number of algorithms, the germline variants make up a substantial fraction of the total calls, showing an association with the number of false positive calls (Figure 2b). Thus germline leakage is, as expected, associated with reduced overall accuracy of mutation calling.

**Figure 2:**
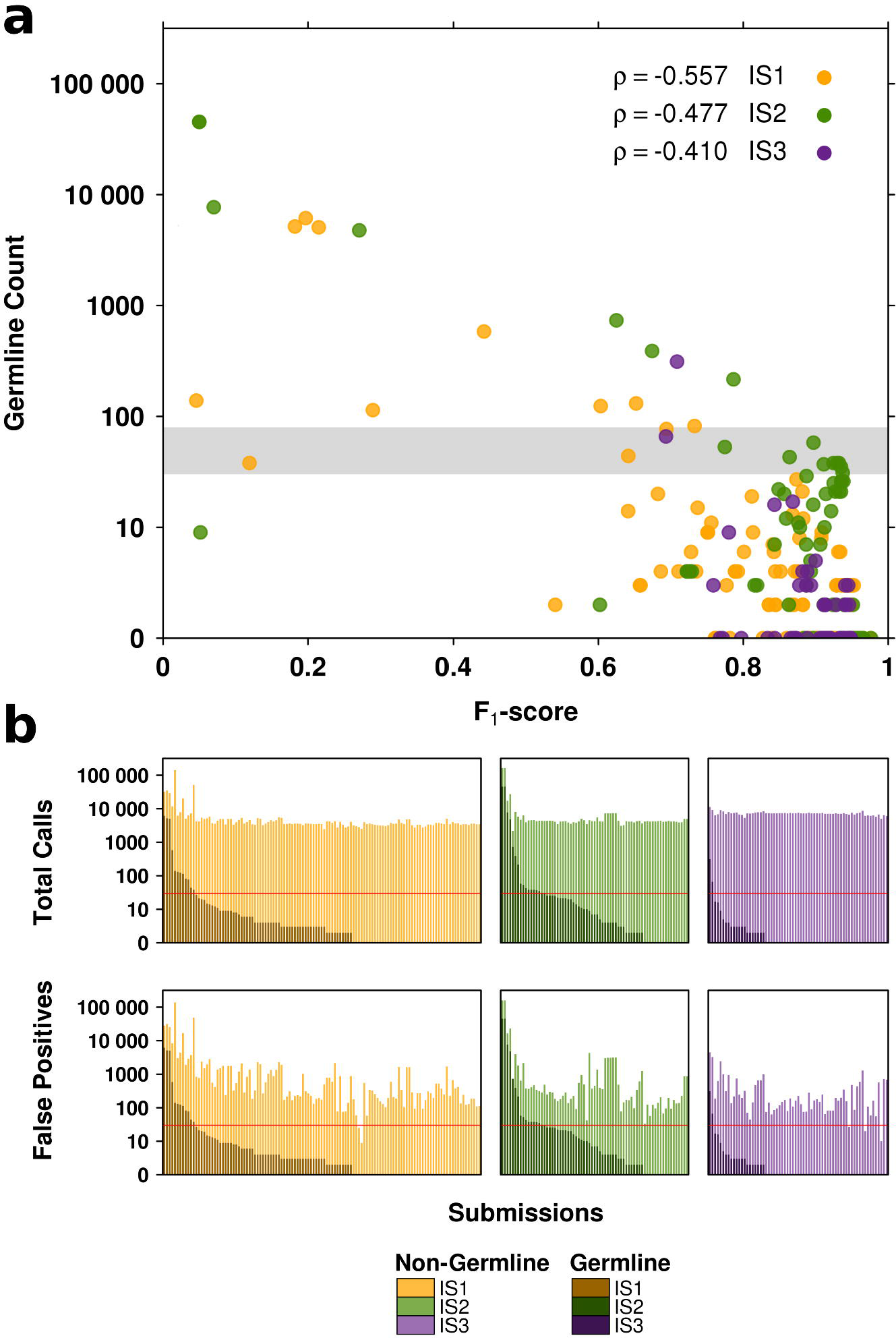
Assessment of somatic SNV prediction accuracy against germline leakage. (**a**) F_1_-scores for each submission are plotted against the germline count (as determined by GermlineFilter). Submissions for different tumours are colour-coded (IS1 = orange, IS2 = green, IS3 = purple). The grey area represents 30-80 counts: the minimum number of independent SNPs required to correctly identify a subject, according to Lin *et al.* [15]. (**b**) Proportions of germline calls as found in total submission calls (upper panel) and in false positive submission calls (lower panel) per tumour. The horizontal red lines indicate the 30 count mark (the lower bound of the 30-80 SNP range mentioned above).

### Quantifying Germline Leakage across Tumours and Between Algorithms

Submissions were further analyzed to determine recurrence of individual germline contaminants across the mutation calling algorithms. For these purposes, only the highest F_1_-score submission from each team was selected, as in the primary report of the somatic SNV data [25]. This was done separately for each tumour, resulting in 15 submissions for IS1, 12 for IS2 and 11 for IS3. A plurality of submissions harboured no germline variants (IS1 = 40.0%; IS2 = 41.7%; IS3 = 45.5%), but there was substantial variability, with one submission containing 43 germline SNPs (**Additional file 2: Supplementary Table 2**).

Individual leaked germline variants varied significantly across algorithms (Figure 3). Of the 85 germline variants leaked in the 12 IS2 submissions (all with an F1 > 0.863), only five were identified more than once. Similarly, of the 23 germline variants leaked in the 11 IS3 submissions, only two were identified more than once. Leaked variants were distributed uniformly across chromosomes. These data suggest that in modern pipelines, germline leakage rates are low and different variants are leaked by different pipelines.

**Figure 3:**
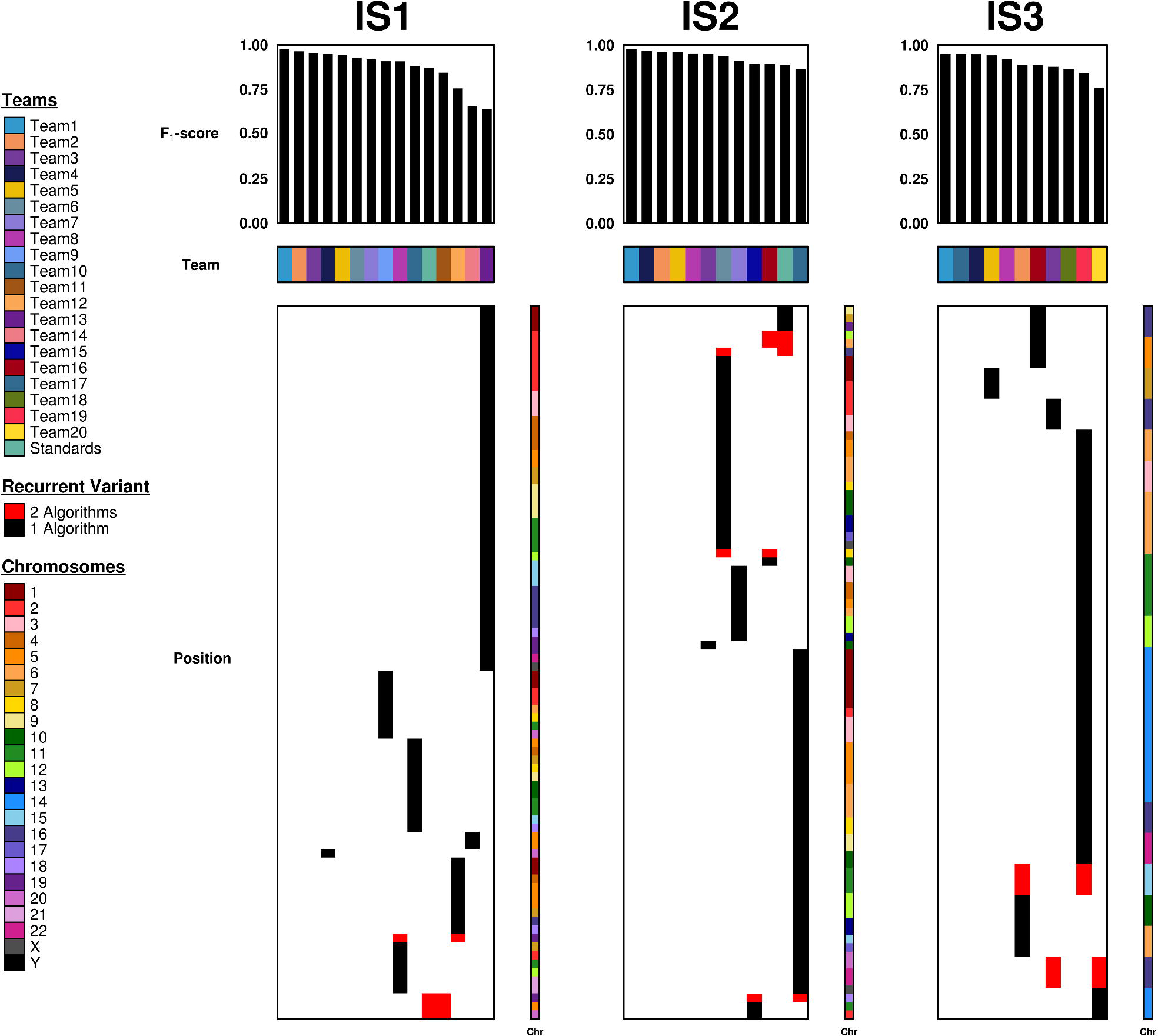
Germline leakage across all tumours (IS1, IS2, IS3) and SNV-calling algorithms. Teams are consistently colour-coded across multiple tumours. Barplots show F_1_-scores from each team’s top-scoring submission. Leaked variants are displayed below with their corresponding chromosomes. Variant bars that overlap horizontally represent recurrent germline leaks.

In addition to the spiked-in mutations, common SNP sites were also analyzed. The Exome AggregationConsortium (ExAC) has produced a library of variant sites seen across 60,706 individuals [27]. These sites represent locations where samples commonly deviate from the reference. Due to the very large number of individuals represented, this set of SNP sites is often used as a filter of possible germline variant sites. ExAC provides ~9.3 million potential common SNP sites, much more than the thousands of spiked-in mutations. The number of false positive calls using ExAC as a filter remained very low (medians: IS1 = 2; IS2 = 3; IS3 = 1.5). As these sites are publicly available and known to be common for SNPs, most modern somatic calling pipelines can directly incorporate this information into their filtering strategy.

## Discussion

Barrier-free access to genomic data can expand its utility, maximizing investments in research funding, enabling citizen-scientists and facilitating collaboration. Strong barriers to access can limit these positive consequences of large investments in dataset generation. Indeed, even when data is made available through protected databases, the processes to gain access can be time-consuming, advantaging labs or institutions that have resources dedicated to gaining and maintaining data-access authorizations. Accessibility can be skewed by variability in the standards, knowledge and impartiality of data access committees that authorize use of controlled data [28-29].

We quantified the amount of leakage in three comprehensively studied tumours used in a crowd-sourced prediction benchmarking challenge. While some submissions showed large amounts of germline leakage, the median submission leaked only one germline SNP, and indeed the top three teams for each tumour leaked none. Given that the SMC-DNA Challenge was run in 2014-2015 and that detection pipelines and the quality of genomic data have improved further since, it appears that modern optimized variant-calling pipelines leak an insignificant number of germline variants on many tumours, well below the 30-80 independent SNP range needed for re-identification [15].

However, several caveats must be evaluated when considering barrier-free access to whole-genome somatic SNV predictions. First, the data we evaluated only included three tumours, and further evaluations on larger numbers with a range of cellularities will be critical to generalize these conclusions. Additionally, while we considered the amount of germline leakage in tumors with different subclonal complexities, we did not investigate whether germline leakage is more likely in genomic regions with specific tumour characteristics (*e.g*. mutational hotspots, trinucleotide context, subclonality, copy number alterations, loss of heterozygosity, *etc*.). On-going work from the ICGC Pan-Cancer Analysis of Whole Genomes (PCAWG) may provide the data necessary to address this. Second, genomic alterations other than nuclear SNVs (*e.g*. germline copy number variants and mitochondrial polymorphisms) may provide information contributing to identifiability. Third, while most individual pipelines leaked few variants, aggregating multiple pipelines could increase the information content: the union of variants across all 12 pipelines from IS2 contain 85 leaked SNPs, potentially providing sufficient information for re-identification [15]. Since ensemble calling generally adopts a ‘majority rules’ approach [30], which would remove most germline variants due to low recurrence, this is most relevant in cases of malicious intent.

Taken together, our findings suggest that germline contamination in somatic SNV calling is relatively rare, and supports additional consideration of barrier-free access to these data. Re-identification risks can be substantially reduced by incorporating automated checks into the data release process, designed to identify germline leakage and remove these prior to data release. GermlineFilter provides a convenient and secure way to monitor leakage by individual algorithms, and may be useful as a front-end to cloud-based SNV databases to quantify and minimize risk in real-time.

## Methods

### Software

GermlineFilter works in an encrypted fashion, allowing its use on a public server. It is executed in two steps (Figure 1). For the first step, performed offline, a VCF file containing germline calls is generated using paired tumour and normal BAM files. The user can additionally specify alternative germline call sets, such as a list of all dbSNP entries, although these would elevate the false-negative rate by removing true somatic mutations. For each germline SNP in the VCF file, we extract the chromosome, position, reference base and alternate base. This information is hashed and written to a file that is then encrypted. It is this file of hashes rather than the actual variants that is then transferred to the server. Next, online somatic VCF filtering is performed. At runtime, the truth germline VCF is decrypted in memory and the somatic VCF undergoes preprocessing and hashing. Finally, an in-memory comparison of hashes is done and the number of matches is returned. GermlineFilter can spawn multiple instances to process multiple germline VCFs for different tumours or multiple somatic VCFs for a single tumour. The user chooses the encryption and hashing protocols, with strong default settings in place to help minimize risks such as hash collisions. An additional feature for local use allows the user to obtain a list of the actual positions of the germline leaks within the somatic VCF. This list can be used to filter out the germline mutations in preparation for publication.

The GermlineFilter software package was written in Python 2.7 and it is supported for Unix and Linux platforms. The encryption and hashing is done using the *PyCrypto* v2.6.1 Python module. The tool currently supports two encryption protocols -- *AES* (default) and *Blowfish*, as well as two hashing protocols -- *SHA512* (default) and *md5*, selected for their security and broad usage. GermlineFilter v1.2 is the stable version and it is available for download at: https://pypi.python.org/pypi/GermlineFilter. Alternatively, it can be installed via pip install GermlineFilter.

### Data

The analysis data was taken from Ewing *et al.* [25] and it consists of the first three publicly available *in silico* datasets from the ICGC-TCGA DREAM Somatic Mutation Calling Challenge and their corresponding SNV submissions from the challenge participants. The truth germline calls were generated using *GATK HaplotypeCaller* v3.3. A description of the synthetic tumour data and a summary of participating teams and their submissions can be found in **Additional file 1: Supplementary Table 1**. All challenge submissions and their scores are listed in **Additional file 2: Supplementary Table 2**.

For each of the 259 submissions we calculated: precision (the fraction of submitted calls that are true somatic SNVs), recall (the fraction of true somatic SNVs that are identified by the caller) and the F_1_-score (the harmonic mean of precision and recall), as previously reported [25]. The F_1_-score was selected to be the accuracy metric as it does not rely on true negative information which, given the nature of somatic variant calling on whole genome sequencing data, would overwhelm alternative scoring metrics such as specificity (the fraction of non-SNV bases that are correctly identified as such by the caller).

Each tumour’s germline calls were encrypted separately using default methods: AES for encryption and SHA512 for hashing. Somatic calls from all challenge submissions were filtered against their corresponding tumour’s encrypted germline calls. For a somatic SNV call to be designated a germline leak, it exactly matched a germline variant at the chromosome, position, reference allele and alternate allele.

The resulting germline leak counts were compared to F_1_-scores using Spearman correlation. The best team submissions per tumour were selected to look at leaked germline variant recurrence across tumours and mutation callers. Best submissions were defined as having the highest F_1_-score.

### Visualization

All data figures were created using custom R scripts executed in the R statistical environment (v3.2.3) using the *BPG* (v5.6.8) package [31].

## List of abbreviations used

BAM: binary alignment map
DREAM: Dialogue on Reverse-Engineering Assessment and Methods
GATK: Genome Analysis Toolkit
HIPAA: Health Information Portability and Accountability Act
ICGC: International Cancer Genome Consortium
NGS: next-generation sequencing
PGP: Personal Genome Project
SMC: Somatic Mutation Calling
SNP: single nucleotide polymorphism
SNV: single nucleotide variant
TCGA: The Cancer Genome Atlas
VCF: variant call format

## Declarations

### Ethics approval and consent to participate

Not applicable

### Consent for publication

Not applicable

### Availability of data and materials

The datasets supporting the conclusions of this article are available on Synapse (syn312572) at: https://www.synapse.org/#!Synapse:syn312572/wiki/61509

And in the Supplementary of Ewing *et al*. [25]. The main GermlineFilter project page is at:https://labs.oicr.on.ca/boutros-lab/software/germlinefilter

And the source-code is freely available at: https://pypi.python.org/pypi/GermlineFilter/1.2

## Authors’ contributions

ADE, AAM, JMS and PCB initiated the project. CC created GermlineFilter and performed validation studies. KE, CC, JCB, TCN, AAM, JMS and PCB created the ICGC-TCGA DREAM Somatic Mutation Calling Challenge. DHS, CC, TNY, KEH and ADE created datasets and analyzed submission data. Research was supervised by AAM, JMS and PCB. The first draft of the manuscript was written by DHS, and approved by all authors.

## Acknowledgements

The authors thank all members of the Boutros lab and all ICGC-TCGA DREAM Somatic Mutation Calling Challenge Participants for their support and thoughtful commentary.

## Additional files

### Additional file 1: Supplementary Table 1

XLS 12.3 kb Tumour information from each tumour challenge (IS 1, IS2, IS3). This includes information on *in silico* tumour construction, composition, and a summary of participating teams and their challenge submissions.

### Additional file 2: Supplementary Table 2

XLS 36.4 kb Contains the following information for every challenge submission: tumour, submission ID, precision, recall, F1-score, and the number of germline variants leaked.

